# LDBlockShow: a fast and convenient tool for visualizing linkage disequilibrium and haplotype blocks based on variant call format files

**DOI:** 10.1101/2020.06.14.151332

**Authors:** Shan-Shan Dong, Wei-Ming He, Jing-Jing Ji, Chi Zhang, Yan Guo, Tie-Lin Yang

## Abstract

The triangular correlation heatmap aiming to visualize the linkage disequilibrium (LD) pattern and haplotype block structure of SNPs is ubiquitous component of population-based genetic studies. However, current tools suffered from the problem of time and memory consuming, and direct calculation from variant call format (VCF) files is not supported. Here we developed LDBlockShow, an open source software, for visualizing LD and haplotype blocks from VCF files. It is time and memory saving. In a test dataset with 100 SNPs from 60,000 subjects, it was at least 429.03 times faster and used only 0.04% – 20.00% of physical memory as compared to other tools. In addition, it could generate figures that simultaneously display additional statistical context (e.g., association *P* values) and genomic region annotations. It can also compress the SVG files with large number of SNPs and support subgroup analysis. This fast and convenient tool would facilitate the visualization of LD and haplotype blocks for geneticists.

## Introduction

Due to genetic linkage, nearby single nucleotide polymorphisms (SNPs) are often highly correlated. In genetic studies, understanding the linkage disequilibrium (LD) pattern is helpful in selecting representative SNP subsets and interpreting other statistical results, such as association *P*-values [1]. However, for multiple SNPs, it can be difficult to interpret results from the summary statistics of pairwise LD measurements since the number of measurements increases rapidly with the number of SNPs. Therefore, the triangular correlation heatmaps aiming to visualize the LD pattern and haplotype block structure of SNPs have become a ubiquitous component of population-based genetic studies since the completion of the international HapMap project.

Haploview [2] and LDheatmap [3] are the most popular tools for visualizing LD heatmaps. Nowadays with files containing large number of individuals and SNPs generated from next generation sequencing (NGS) data, both tools suffered from the problem of time and memory consuming. In addition, in order to display regional association statistics or genomic annotation results in the context of LD, researchers should manually align two disjoint figures. For example, previous studies [4-6] have merged the association statistics, recombination rate, or genomic region annotation results with the LD plots generated by Haploview, which is inconvenient. Moreover, with large number of SNPs, the produced SVG or PDF vector diagram might be too large to open in personal computers. For example, with 1,000 SNPs, a total of 500,000 grids (1000 × 1000 / 2) will be generated and the plot file would reach to tens of MB. Haploview supports the output of PNG file, but it might suffer from the problem of low resolution. Besides, with variant call format (VCF) [7] files generated from NGS data analysis, researchers should convert the VCF files to “PED” file format first, and then use Haploview or LDheatmap to get the LD plot.

To address the above mentioned problems, here we present a software, LDBlockShow, to allow biologists to generate LD and haplotype maps quickly and directly from VCF files. LDBlockShow supports the generation of LD heatmap and regional association statistics or genomic annotation results simultaneously. LDBlockShow can also compress the SVG files with large number of SNPs. This fast and convenient tool would facilitate the visualization of LD and haplotype blocks for geneticists.

## Results

### Overview of LDBlockShow workflow

LDBlockShow consists of two units, the data processing unit and the plot unit (Fig. 1). The data processing unit takes compressed or uncompressed VCF files as input. Users can also input files in PLINK [8] or genotype format with the option of “*-InPlink*” and “*-InGenotype*”, respectively. Next, subgroup samples can be selected (“-*Subgroup*” flag), and SNPs with minor allele frequency (MAF) of less than 0.05, or missing sample rate of over 0.25, or heterozygosis ratio of over 0.9 will be filtered. Custom criteria can be defined with *“-MAF”, “-Miss”*, and *“-Het”*. The genotype of each individual will be stored in specific data structure to facilitate pairwise LD statistics calculation. Using the calculated LD measurement statistics, the plot unit generates the final LD heatmap. Specifically, users can generate the LD plot combined with additional statistical context or genomic region annotations simultaneously. With large number of SNPs, LDBlockShow will compress the output SVG file.

**Figure 1.**
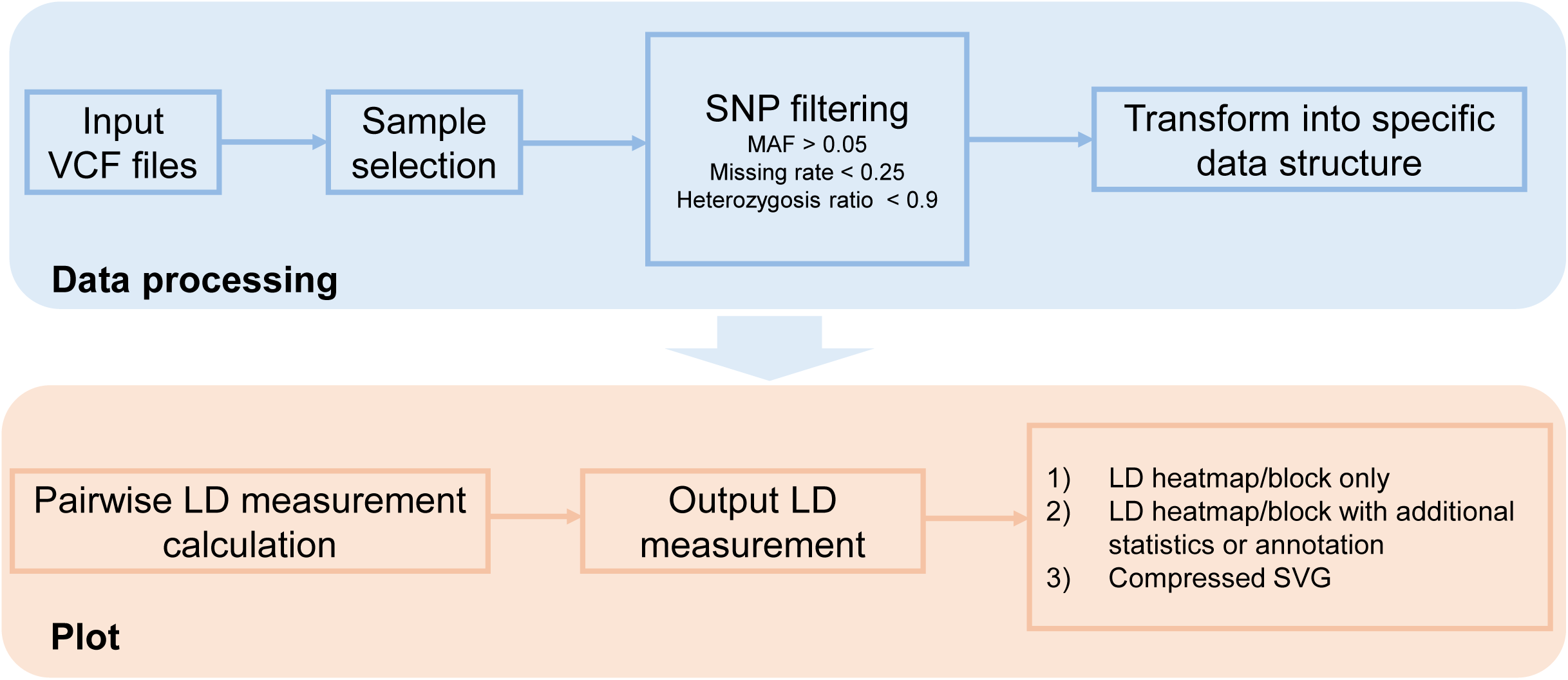
The LDBlockShow workflow. It consists of two units, the data processing unit and the plot unit. The data processing unit include sample selection, SNP filtering and data structure transformation. The plot unit will generate the LD heatmap with calculated LD measurement statistics.

### LDBlockShow is time and memory saving

We examined the computing time and memory requirement of LDBlockShow, Haploview [2], and LDheatmap [3] using genotype data from the UK Biobank population (all SNPs on chromosome 22). As shown in Figure 2, with a fix SNP number of 100 and sample size ranging from 2,000 to 60,000, LDBlockShow was much faster than the other two tools. For example, with the sample size of 60,000, it took LDheatmap and Haploview 98.68 and 123.87 minutes to generate the LD plot. In contrast, it took LDBlockShow only 0.23 minutes to analyze the same dataset, representing 429.03-, and 538.57-fold speed gain over these two methods. In addition, LDBlockShow only required a small amount of physical memory. For example, with the sample size of 60,000, LDheatmap and Haploview required 0.15, and 75.00 GB memory for analyzing 100 SNPs, respectively. In contrast, LDBlockShow used only 0.03 GB, which is 20.00%, and 0.04% of that required by the two methods. With a fixed sample size of 2,000 and SNP number ranged from 100 to 1,200, LDBlockShow showed similar performance (Fig. 2C and Fig.2D). For example, with 1,200 SNPs, it took LDheatmap and Haploview 529.13 and 1289.88 minutes to generate the LD plot. In contrast, it took LDBlockShow only 0.23 minutes to analyze the dataset, representing 2300.57-, and 5608.17-fold speed gain over these two methods. In the same dataset, LDheatmap and Haploview required 0.28, and 11.00 GB memory, respectively. LDBlockShow used 0.10 GB, which is 55.56%, and 0.91% of that required by the two methods.

**Figure 2.**
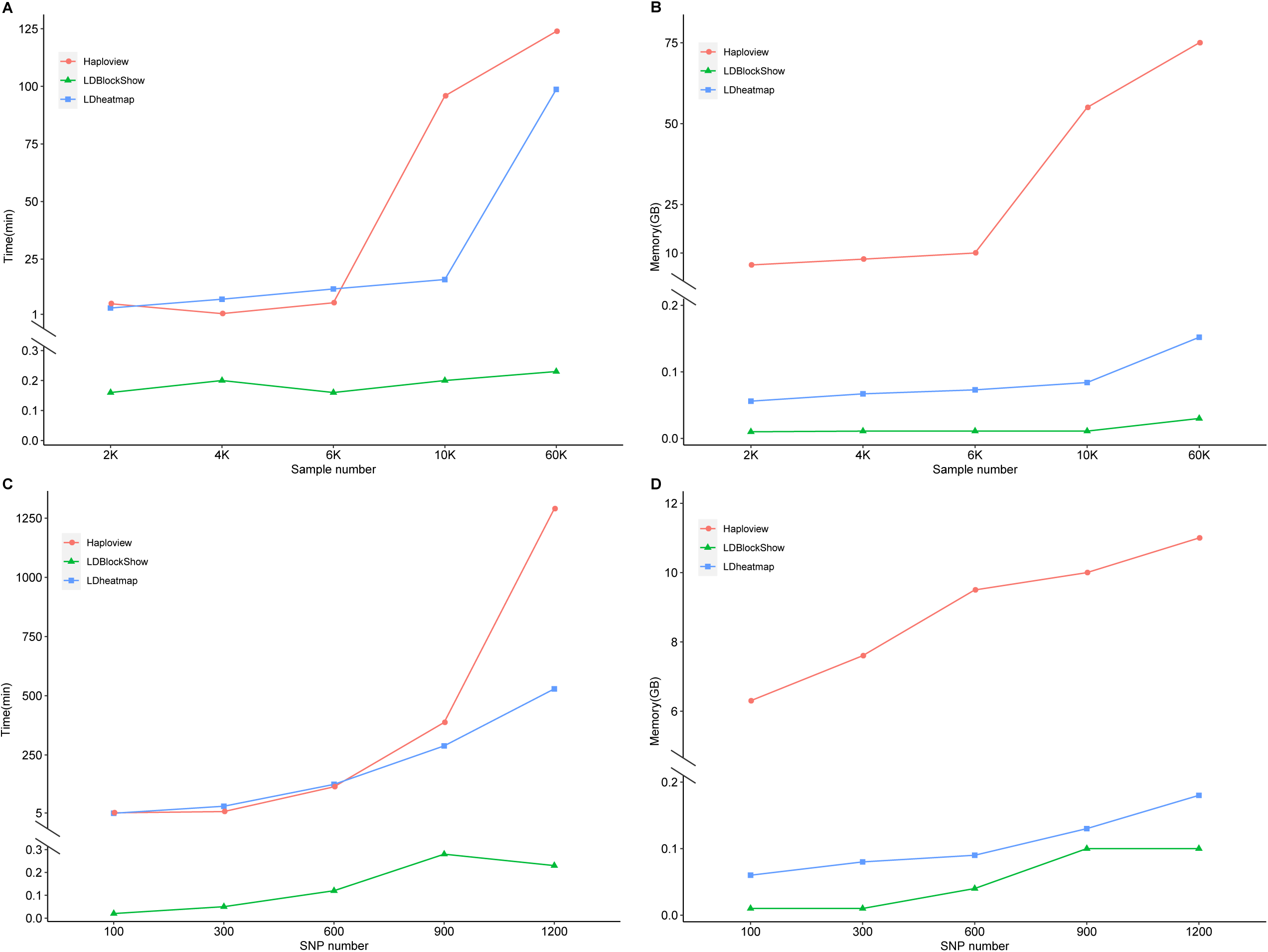
Comparison of computing cost for LDBlockShow, LDheatmap and Haploview. CPU time (A) and memory cost (B) for different methods are shown with a fix SNP number of 100 and sample size ranging from 2,000 to 60,000. CPU time (C) and memory cost (D) for different methods are shown with a fixed sample size of 2,000 and SNP number ranged from 100 to 1,200. Computation is performed with one thread of an Intel Xeon CPU E5-2630 v4.

### LDBlockShow is convenient in combing additional statistics or genomic annotations

LDBlockShow can generate the plots of LD heatmap and additional statistical context (provided with the “-*InGWAS*” flag) or genomic annotation results (provided with the “-*InGFF*” flag) simultaneously. An example output plot of LDBlockShow is shown in Fig. 3.

**Figure 3.**
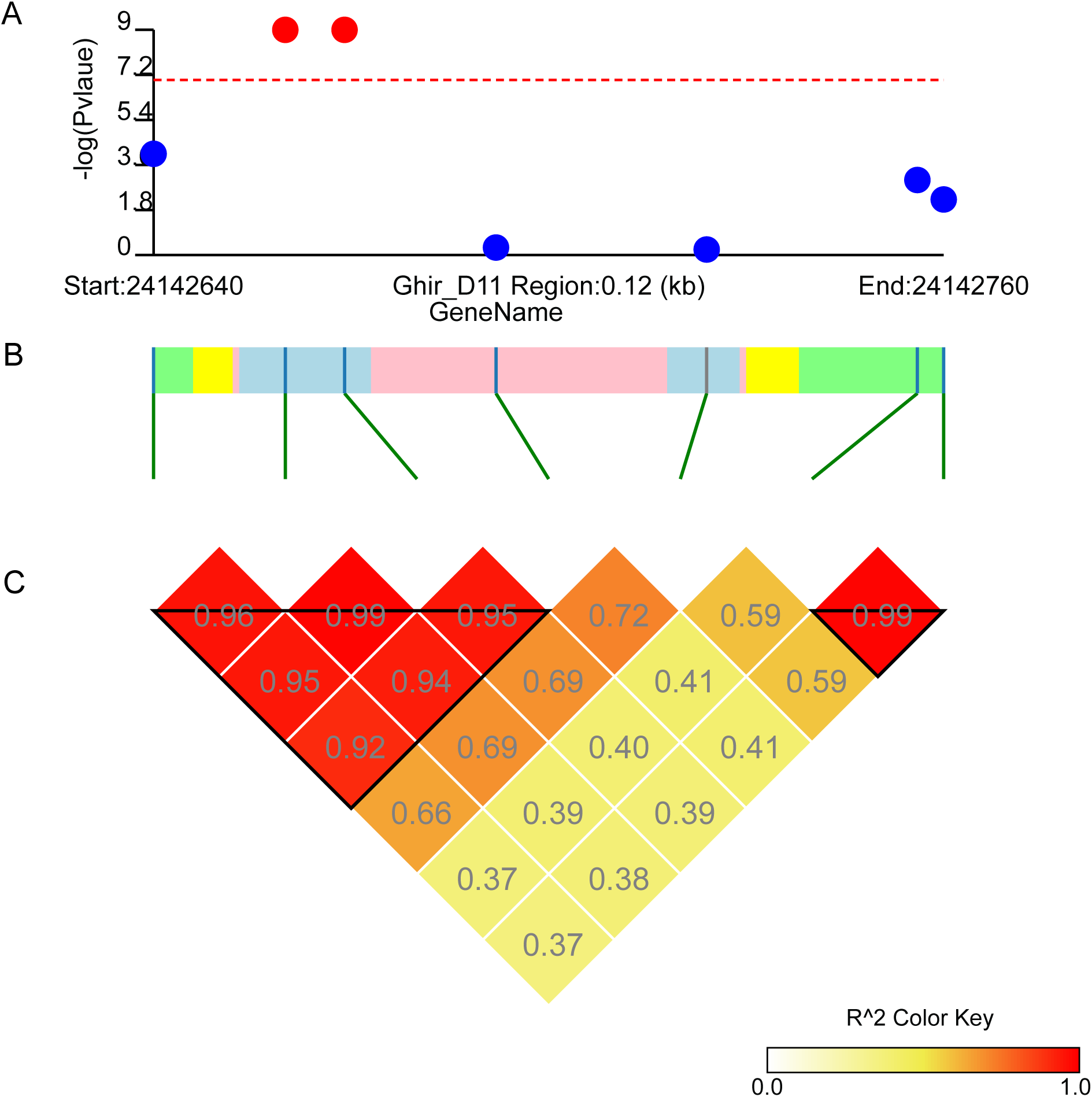
An example output figure of LDBlockShow. A. The association statistics. In addition to the association statics, users can choose other statistical measurements for SNPs to display. B. The genomic region annotation results. By default, CDS, intron and UTR are shown in light blue, pink and yellow, respectively. Other genomic regions are shown in green. Colors can be user-defined. C. The LD heatmap. Colors can be user-defined. The LD measurement values is showed with the flag of “*-ShowRR*”.

### LDBlockShow support the compression of SVG files with large number of SNPs

In addition to converting SVG to PNG file (“-*OutPng*” flag), we also offered another option to compress the SVG file. With SNP number over 50 (cutoff can be defined with the “*-MerMinSNPNum*” flag), the SVG file can be automatically compressed with small number of color gradients. Adjacent grids with the same color will be merged into a single grid. In a test dataset of 1,000 SNPs, under the condition of no compression, the sizes of the heatmap plot files generated by, Haploview, and LDheatmap were 26, 91, and 23 MB, respectively. After compression with LDBlockShow, the above 26MB SVG will be compressed to 8.6 MB (33.08% of the original size). The overall comparison of LDBlockShow, Haploview, and LDheatmap is shown in Table 1.

**Table 1.** Comparison of LDBlockShow with other tools

| **Performance** | **LDBlockShow** | **Haploview** | **LDheatmap** |
| --- | --- | --- | --- |
| ***Input*** |  |  |  |
| Calculation from VCF files directly | √ | × | × |
| Support subgroup analysis | √ | × | × |
| ***Output*** |  |  |  |
| Visualize additional statistics or genomic annotation simultaneously | √ | × | × |
| Compressed SVG | √ | × | × |
| PNG file | √ | √ | × |
| Block region | √ | √ | × |
| LD measurement | $D'/r^2$ | $D'/r^2$ | $r^2$ |

## Discussion

In this study, we developed LDBlockShow, for visualizing LD and haplotype blocks based on VCF files. Compared with current tools, LDBlockShow has the following advantages: Firstly, it is time and memory saving and supporting analyses directly from VCF files. With the advances of NGS, genomic data for large scale populations have been generated gradually. For example, the numbers of human exomes and whole genomes of the genome aggregation database (gnomAD) consortium have reached 125,748, and 15,708, respectively [9]. Therefore, LDBlockShow can offer help for NGS researchers in a time and resource efficient manner. Secondly, LDBlockShow also complements the common triangular correlation heatmaps by providing additional statistical context and genomic region annotations. For example, with association analysis statistics, users can easily locate the SNPs with the most significant association signal, which is especially useful for genomic fine-mapping studies. Thirdly, with large number of SNPs, LDBlockShow could compress the original vector diagram to about 33% of the original file size, facilitating visualization in personal computers. In addition, subgroup analysis is supported by LDBlockShow, which is convenient for users to compare the LD patterns in different subgroups.

## Conclusion

In conclusion, LDBlockShow is a fast and convenient tool for visualizing LD and haplotype blocks based on VCF files. It supports the generation of LD heatmap and regional association statistics or genomic annotation results simultaneously. LDBlockShow can also compress the SVG files with large number of SNPs and support subgroup analysis.

## Methods

### Data structure

LDBlockShow is implemented in C++ with Open MIC license for Linux/Unix and Mac operating system. The genotype of each individual will be stored with the following data structure to facilitate the fast pairwise LD statistics calculation:

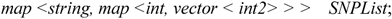

### LD measurement

Pairwise LD measurements D’ and r^2^ were calculated according to previously reported formulas [10, 11]. That’s, suppose allele A occurs with frequency *PA* at one locus and allele B occurs with frequency *PB* at another locus, then we have the D’:

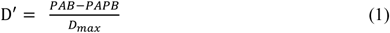

where

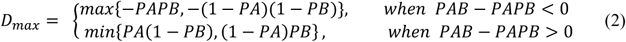

and the r^2^:

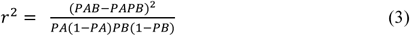

Users can choose to display r^2^ or D’ in the heatmap with the option of “*-SeleVar”*.

### LD blocks

LDBlockShow supports the definition of blocks in four different ways. By default, PLINK (Version 1.9, www.cog-genomics.org/plink/1.9/) [8] will be called to generate the block defined by Gabriel *et al*. [12]. The solid spine of LD [2] method is also supported. Users can also define their own cutoff of r^2^ and D’ for blocks with the option of “*-BlockCut*” or supply their own block region definition with the option of “*-FixBlock*”.

### Code availability

LDBlockShow is freely available for non-commercial research institutions. Details can be obtained from https://github.com/BGI-shenzhen/LDBlockShow.

## Acknowledgement

We thank the LDBlockShow community members that have offered great suggestions for improving the software. This research has been conducted using the UK Biobank Resource under Application Number 46387.

## Authors’ contributions

T.-L.Y., W.-M.H., and S.-S.D. conceived and designed the study. S.-S.D. wrote the paper. W.-M.H. implemented the software with the help of S.-S.D., J.-J.J, and C.Z. Y.G. edited the paper. The authors read and approved the final manuscript.

## Ethics approval and consent to participate

Not applicable.

## Consent for publication

Not applicable.

